# Contributions of β-lactamase substrate specificity and outer membrane permeability to the antibiotic sheltering of β-lactam-susceptible bacteria

**DOI:** 10.1101/2025.04.22.649001

**Authors:** Montserrat Mora-Ochomogo, Mitchell A. Jeffs, Josephine L. Liu, Christopher T. Lohans

## Abstract

The use of β-lactam antibiotics is threatened by antibiotic resistant bacteria that produce β-lactamases. These enzymes not only protect the bacteria that produce them but also shelter other bacteria in the same environment that would otherwise be susceptible. While this phenomenon is of clinical significance, many of the factors that contribute to β-lactamase-mediated antibiotic sheltering have not been well-studied. We report the development of a luminescence assay to directly monitor the survival of β-lactam-susceptible bacteria in the presence of β-lactamase-producing bacteria and β-lactam antibiotics. This method provides a rapid and scalable means of quantifying antibiotic sheltering in mixed microbial populations. We applied this assay to investigate the contributions of several factors to sheltering, including the class of β-lactam, the substrate specificity of the β-lactamase, and the cell wall permeability of the β-lactamase-producing bacterium. Our results show that the extent of sheltering that occurs depends greatly on the particular combination of β-lactam and β-lactamase, and also on the ability of a β-lactamase to access its β-lactam substrate.

## Introduction

Microbes, including the trillions that live in the human body, exist in complex environments which can contain a multitude of different strains and species.^1^ While many members of the human microbiota are beneficial to their host, the growth of pathogenic microbes in the body can lead to infections. Furthermore, some infections result from the presence of multiple different pathogenic microorganisms.^2^ These polymicrobial infections are frequently associated with a number of different health conditions, including cystic fibrosis, chronic obstructive pulmonary disease, wound infections, and dental cavities.^2–7^ The co-occurrence of multiple pathogens has been shown to impact microbial processes such as colonization and virulence, and can complicate antimicrobial therapy.^5,8^ Studies have shown that the ability of an antibiotic to target a bacterial pathogen can be influenced greatly by other bacteria present in the same environment.^9–11^

Bacterial infections are commonly treated with β-lactams, a group which represents more than 50% of the antibiotics prescribed worldwide.^12–14^ The members of this group, including the penicillins, cephalosporins, and carbapenems, contain a characteristic four-membered β-lactam ring which confers them with bactericidal activity. β-lactams target bacterial penicillin-binding proteins (PBPs), disrupting cell wall synthesis by inhibiting the last steps of peptidoglycan formation, ultimately leading to cell lysis.^14^ When used to target Gram-negative bacteria, β-lactams must cross the outer membrane (OM) to reach the periplasmic space where PBPs are located (Figure 1A).

**Figure 1.**
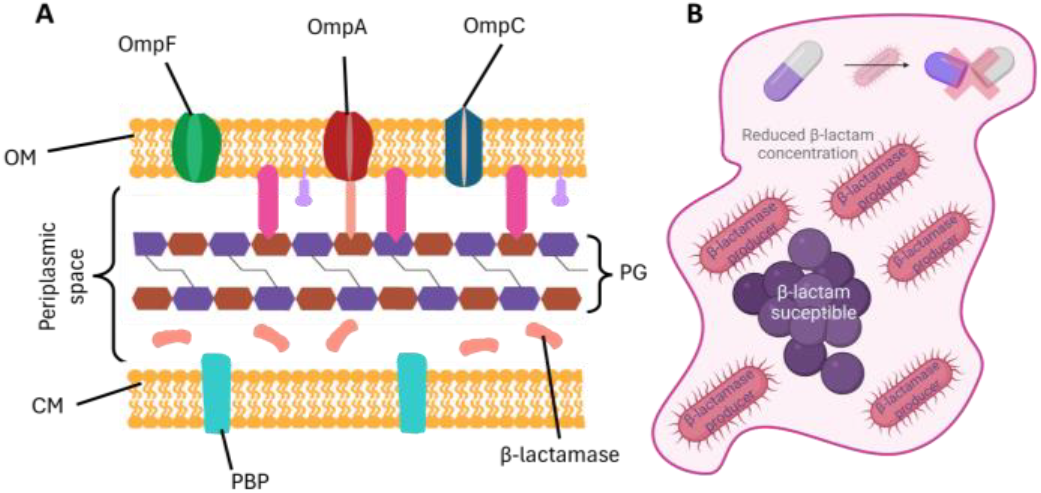
**A.** Overview of the major components in the Gram-negative cell envelope. β-lactams target penicillin-binding proteins located in the periplasm. Outer membrane proteins such as OmpF and OmpC allow β-lactams to enter the periplasm, and proteins Lpp and NlpI play major roles in outer membrane stability. β-lactamase enzymes in the periplasm degrade β-lactams. CM: cytoplasmic membrane; OM: outer membrane; PBP: penicillin-binding protein; PG: peptidoglycan. B. Schematic representation of antibiotic sheltering. β-lactamase-producing bacteria (red) can protect bacteria that are susceptible to β-lactams (purple) by decreasing the local concentration of β-lactam antibiotic present in the environment. Created in BioRender. (2025). https://BioRender.com/el5zavf

Several different resistance mechanisms can protect bacteria against β-lactams, and the production of β-lactamases is especially prevalent among Gram-negative pathogens.^15–17^ β-lactamase enzymes hydrolytically inactivate β-lactam antibiotics, preventing them from targeting PBPs and disrupting peptidoglycan synthesis. More than 8,000 β-lactamase variants have been identified to date,^18^ and the members of this group vary greatly in terms of which β-lactams they can degrade. Penicillinases like TEM-1 are principally active against penicillins and some cephalosporins, while extended-spectrum β-lactamases (ESBLs) like CTX-M-15 and SHV-2 can also degrade later generation cephalosporins.^19,20^ Carbapenemases such as NDM-1, IMP-1, and KPC-2 are a significant concern as they hydrolyze carbapenems, a group of β-lactams often reserved as treatments of last resort for the management of antibiotic-resistant infections.^21,22^

While the substrate specificity of a β-lactamase is determined by the structure of its active site and catalytic mechanism, β-lactamase activity is also impacted by the surface layers of the bacterium that produces it. As β-lactamases are primarily located in the periplasmic space in Gram-negative bacteria (Figure 1A), β-lactam degradation in the periplasm first requires the antibiotic to cross the OM. The entry of many β-lactams is dependent on porins, protein channels in the OM that facilitate the entry of small molecules into the periplasm.^23,24^ Hence, changes to porins and other outer membrane proteins impact the rate of β-lactam degradation.

The production of β-lactamases by pathogenic bacteria can contribute to treatment failure when β-lactam antibiotics are used to treat polymicrobial infections.^12,25^ This likely arises, at least in part, from the protection that β-lactamase-producing bacteria provide to β-lactam-susceptible bacterial populations (Figure 1B).^26–34^ By depleting the amount of extracellular β-lactam that is present, resistant bacteria can help susceptible bacteria survive antibiotic exposure.^35,36^ This phenomenon has been the subject of many recent studies, and has often been referred to as antibiotic sheltering, bacterial cheating, collective resistance, or group beneficial traits.^26,35,37–41^

Although antibiotic sheltering is relevant in the context of the treatment of bacterial infections, the majority of antibiotic susceptibility testing (in both academic and clinical contexts) uses bacterial monocultures. Indeed, the study of sheltering can be challenging, requiring the quantification of susceptible bacteria within a more complex population. To date, many studies in this area have employed plate counting,^27,42,43^ dual flask experiments,^26,41^ and growth cultures paired with mathematical models.^9,38,40^ However, these methods can be labour-intensive, and it can be challenging to differentiate between resistant and susceptible cells, particularly for phenotypically similar strains. Furthermore, some methods use a relatively complex experimental set up with continuous supplementation of components.^39,44^ These drawbacks can complicate the use of such methods for the investigation of the factors that contribute to antibiotic sheltering.

We report the application of a luminescent *Escherichia coli* reporter strain to characterize the antibiotic sheltering provided by β-lactamase-producing bacteria. Sheltering assays using this reporter are fast and versatile, allowing for factors such as β-lactamase identity and cell wall permeability to be quantitatively investigated. We observed that the extent of sheltering is largely determined by the substrate specificity of the β-lactamase being produced, and was closely related to the kinetic rates of β-lactam degradation. OM permeability of the β-lactamase-producing strain also had a major impact on sheltering, and *E. coli* mutants lacking certain porins offered far less protection. Our methods and results will support future studies in this area which may inform on how the treatment of bacterial infections depends on the resistance mechanisms associated with other microbes present in the body.

## Results and discussion

To investigate factors that determine the antibiotic sheltering provided by β-lactamase-producing bacteria, we prepared a luminescent β-lactam-susceptible *E. coli* BW25113 reporter strain transformed with the pJ23100LUX plasmid. This strain constitutively expresses the *lux* operon, providing a rapid means for evaluating its growth and survival. Initial validation experiments showed that luminescence intensity is related to the number of reporter cells present, and luminescence and OD600 measurements followed similar trends during the growth of *E. coli* BW25113 pJ23100LUX (Figure S2). Addition of the β-lactam antibiotics meropenem or imipenem to the culture led to a decrease in luminescence over time reflecting the killing of the reporter strain (Figure S3).

We next tested whether *E. coli* cells producing the β-lactamase KPC-2 can rescue the luminescent *E. coli* reporter strain from β-lactam antibiotics. A sample containing the reporter strain, a KPC-2-producing strain, and the carbapenem imipenem emitted a strong luminescent signal, while a sample in which the KPC-2-producing strain was substituted with a non-β-lactamase-producing strain had very low luminescence (Figure 2A). These measurements are consistent with the KPC-2-producing strain sheltering the reporter strain by degrading imipenem. Testing multiple β-lactam concentrations in parallel allowed for dose-response curves to be obtained, as shown in sheltering experiments with the carbapenem meropenem and an *E. coli* strain that produces the β-lactamase NDM-1 (Figure 2B). These dose-response data allowed for the determination of the effective concentration at which the growth of the β-lactam-susceptible strain [measured in relative luminescence units (RLU)] is inhibited by half (i.e., an EC_50_ value). Different cell densities of the NDM-1 producing strain were tested for their ability to shelter the reporter strain from a range of meropenem concentrations. As expected, the EC_50_ values showed that higher numbers of NDM-1-producing cells led to increased levels of sheltering. β-lactamase expression levels were also shown to impact the level of sheltering that occurs, with *E. coli* cells transformed with the high copy pHSG298-NDM-1 plasmid sheltering the reporter strain from meropenem at a level approximately three times greater than seen for *E. coli* transformed with the lower copy pACYC184-NDM-1 (Figure S4).

**Figure 2.**
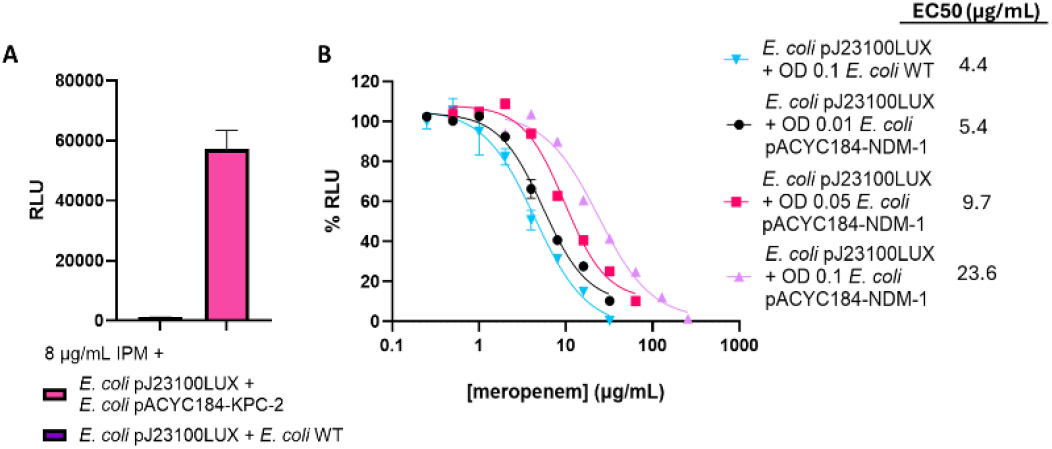
Validation of the luminescence-based sheltering assay. A. Luminescence measurements obtained from the sheltering assay when the reporter strain was treated with 8 µg/mL of imipenem in the presence and absence of KPC-2-producing bacteria. n = 4. B. Dose-response curves showing the impact of different meropenem concentrations on the luminescence of the reporter strain in the presence of different cell densities of NDM-1-producing *E. coli*. 95% confidence intervals are reported in Table S1. n = 3. RLU: relative luminescence units; IPM: imipenem; WT: wild-type. Error bars represent standard deviations.

Following these initial experiments, we carried out assays evaluating the level of sheltering provided by a panel of *E. coli* strains producing different clinically relevant β-lactamases against selected β-lactam antibiotics. Of the β-lactamases tested, NDM-1, KPC-2, IMP-1, and OXA-48 are carbapenemases, ^21,45,46^ while TEM-116 is a penicillinase and CTX-M-15 is an ESBL.^47^ EC_50_ values were determined for selected combinations of β-lactamases and β-lactams, with a focus on sheltering involving the carbapenems meropenem and imipenem. While β-lactamases with carbapenemase activity can degrade carbapenems, these antibiotics are not efficiently degraded by penicillinases and ESBLs.^48,49^

Most of the carbapenemase-producing *E. coli* strains dramatically improved the survival of the reporter strain in the presence of meropenem, with the extent of sheltering varying according to the identity of the carbapenemase (Figure 3, Table S1). NDM-1-producing *E. coli* increased the EC_50_ by 5-fold compared to non-β-lactamase-producing *E. coli* (Figure 3A), while *E. coli* producing KPC-2 and IMP-1 increased the EC_50_ by 8- and 10-fold, respectively (Figures 3B, 3C). While OXA-48-producing *E. coli* provided almost no sheltering, with just a 1.5-fold increase in EC_50_ (Figure 3D), this is consistent with the relatively low catalytic activity of OXA-48 against carbapenems.^50,51^ As expected, *E. coli* producing the penicillinase TEM-116 did not alter the EC_50_ when compared to the unsheltered control (Figure 3E).

**Figure 3.**
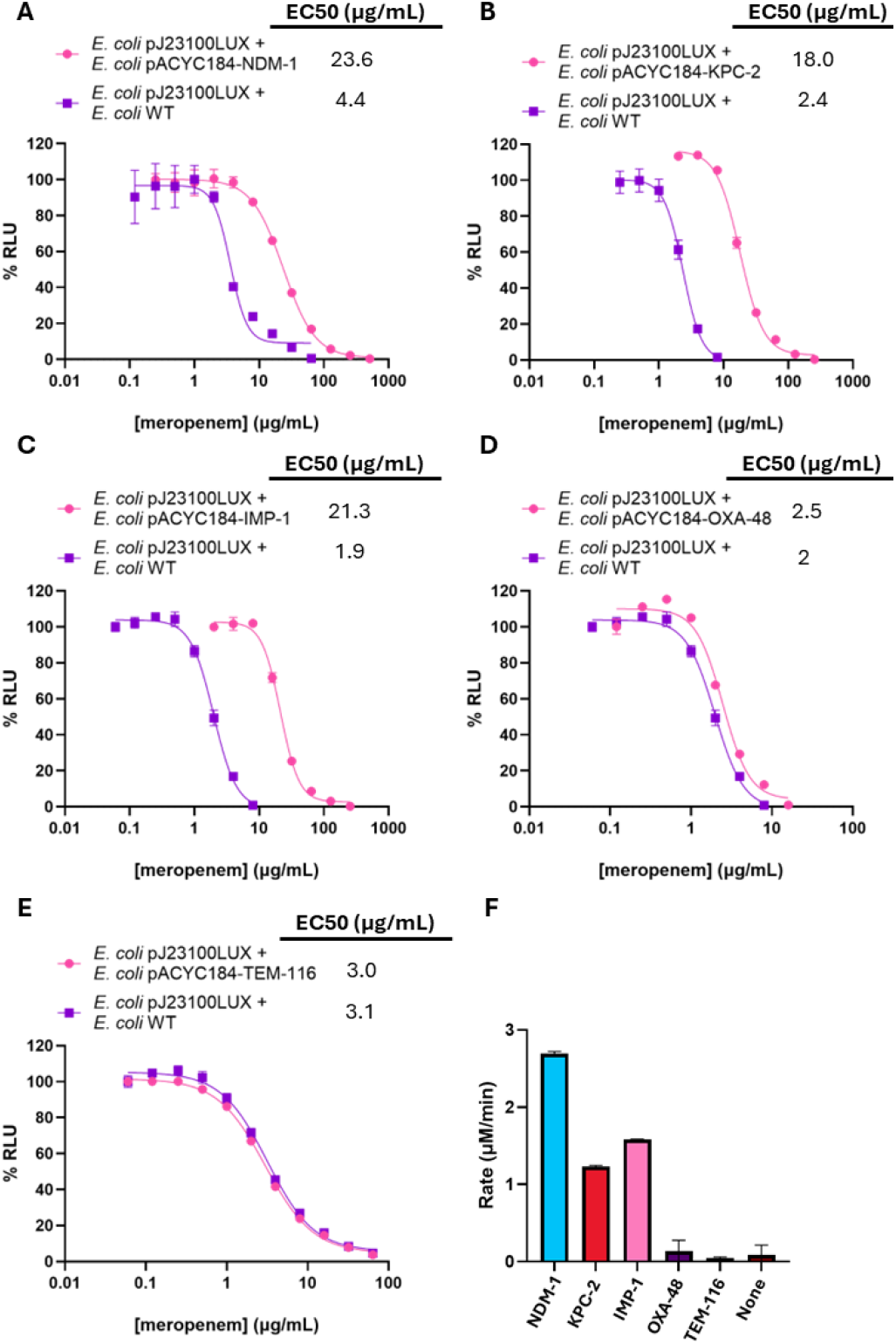
Sheltering by β-lactamase-producing *E. coli* against the carbapenem meropenem. Dose-response curves used to determine EC_50_ values for the luminescent reporter strain in the presence of meropenem and *E. coli* strains producing A. NDM-1, KPC-2, C. IMP-1, D. OXA-48, and E. TEM-116. F. Initial velocities for meropenem (167 μM) hydrolysis by β-lactamase-producing cells as determined by UV-Vis spectrophotometry. 95% confidence intervals are reported in Table S1. % RLU = percent normalized relative luminescence units. n = 4. Error bars represent standard deviations.

We related the sheltering assay results to β-lactamase activity by carrying out UV-Vis spectroscopic assays measuring the degradation of meropenem by the β-lactamase-producing bacteria tested (Figure 3F). The trends observed in the EC_50_ values were consistent with the initial velocities determined in these kinetic assays. Of the strains tested, the NDM-1-producing *E. coli* degraded meropenem most quickly and provided the greatest level of sheltering. *E. coli* strains producing KPC-2 and IMP-1 degraded meropenem more slowly, and provided more moderate levels of sheltering. The very low levels of meropenem hydrolysis catalyzed by *E. coli* producing OXA-48 and TEM-116 was consistent with the poor levels of sheltering observed.

Carbapenemases vary in terms of how efficiently they degrade different carbapenems. Exploring this selectivity, we carried out sheltering assays with the carbapenem imipenem in place of meropenem. Similar to what was seen with meropenem, *E. coli* strains producing the carbapenemases NDM-1 and KPC-2 sheltered the susceptible reporter strain from imipenem, increasing the EC_50_ values by more than 6-fold compared to a non-β-lactamase producing control (Figures 4A, 4B, Table S1). This observation is consistent with reports that KPC-2 degrades imipenem more efficiently than meropenem.^52^ Although the OXA-48-producing strain offered less protection (Figure 4C, Table S1), the fold increase in EC_50_ for imipenem was greater than what was seen for meropenem, as expected based on the kinetic properties of OXA-48.^53^ The trends observed in the imipenem sheltering assays align with the initial velocities of imipenem hydrolysis for NDM-1-, KPC-2- and OXA-48-producing *E. coli* strains as measured by UV-Vis spectrophotometry (Figure 4D).

**Figure 4.**
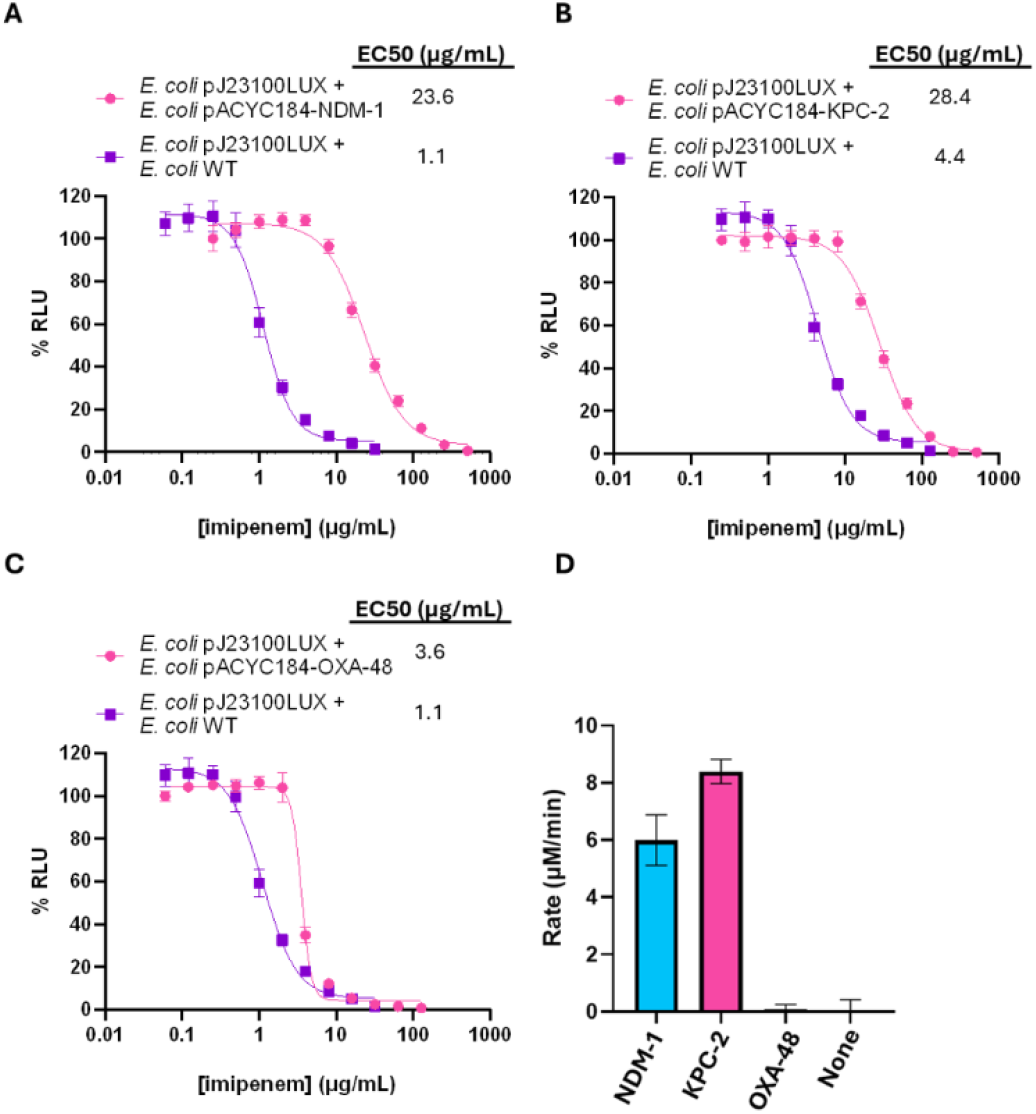
Sheltering by carbapenemase-producing *E. coli* against the carbapenem imipenem. Dose-response curves used to determine the EC_50_ values for the luminescent reporter strain in the presence of imipenem and *E. coli* cells producing A. NDM-1, B. KPC-2, and C. OXA-48. D. Initial velocities for imipenem (214 μM) hydrolysis by β-lactamase-producing cells as determined by UV-Vis spectrophotometry. 95% confidence intervals are reported in Table S1. % RLU = percent normalized relative luminescence units. n = 4. Error bars represent standard deviations.

β-lactam antibiotics are often prescribed in combination with β-lactamase inhibitors as a countermeasure against β-lactamase-producing bacteria. There are several clinically approved inhibitors which target serine β-lactamases (SBLs) (e.g., KPC-2), preventing these enzymes from degrading β-lactam antibiotics.^13^ To test whether the presence of a β-lactamase inhibitor impacts the level of sheltering that occurs, we carried out a sheltering assay with KPC-2-producing *E. coli*, the β-lactamase inhibitor avibactam,^54^ and meropenem (Figure 5A). The addition of 4 μg/mL avibactam prevented measurable sheltering from occurring, lowering the EC_50_ for a sample containing KPC-2-producing *E. coli* to the level observed for a non-β-lactamase-producing control (Figure 5A, Table S1).

**Figure 5.**
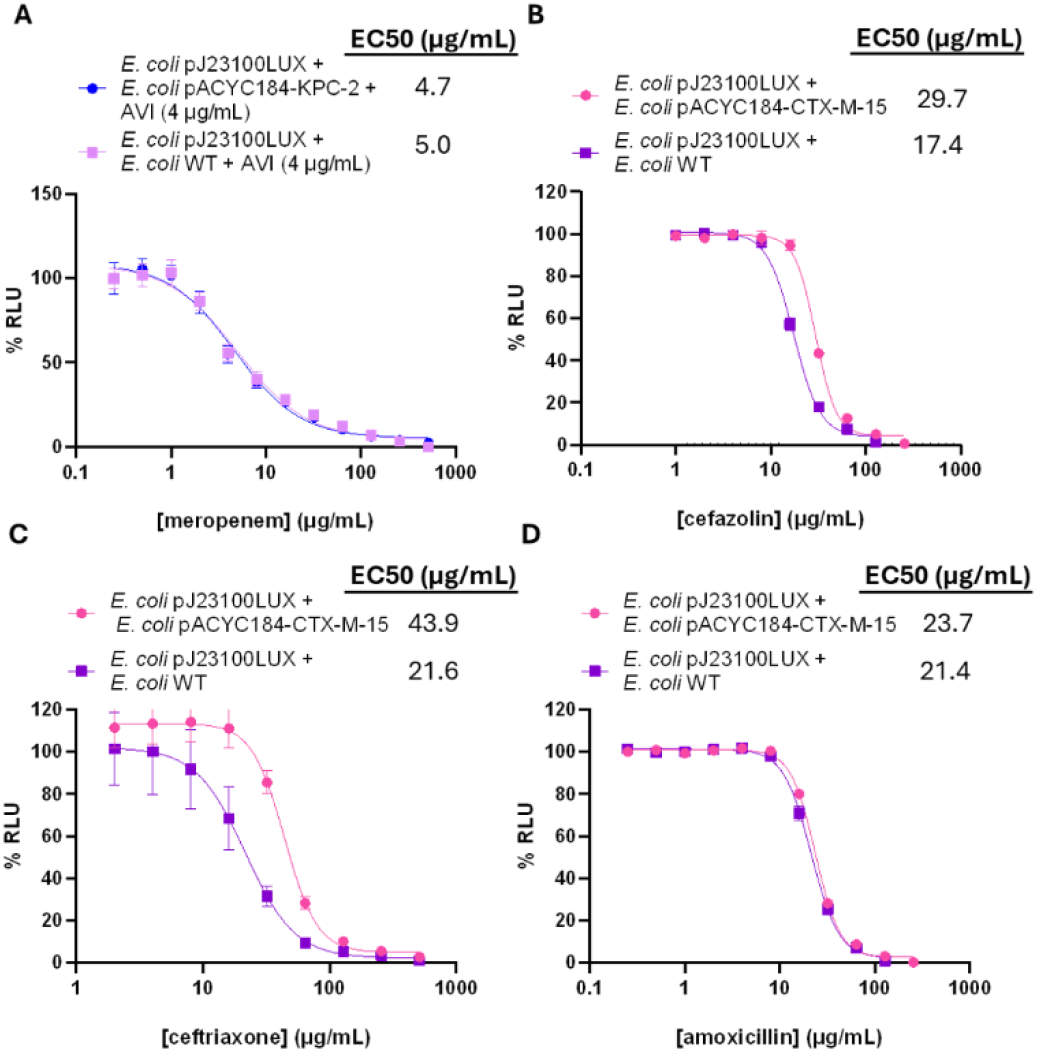
Sheltering experiments with β-lactamase inhibitors and other β-lactam classes. A. Dose-response curves showing that the EC_50_ value for KPC-2-producing *E. coli* treated with meropenem and the β-lactamase inhibitor avibactam (AVI) is similar to that obtained for non-β-lactamase-producing *E. coli*. Dose-response curves showing the EC_50_ values for the reporter strain in the presence of CTX-M-15-producing *E. coli* cells and B. cefazolin, C. ceftriaxone, and D. amoxicillin. 95% confidence intervals are reported in Table S1. % RLU = normalized percent relative luminescence units. n = 4. Error bars represent standard deviations.

Although our studies were primarily focused on sheltering involving carbapenems and carbapenemase-producing bacteria, we tested whether the assay could be applied to investigate antibiotic sheltering involving other types of β-lactams. Bacteria that produce ESBLs are a major clinical concern because of the ability of these enzymes to hydrolyse extended-spectrum cephalosporins such as ceftriaxone.^55,56^ We tested the sheltering offered by *E. coli* producing the ESBL CTX-M-15 against cefazolin (first generation cephalosporin) and ceftriaxone (third generation cephalosporin) (Figures 5B, 5C, Table S1). Comparable levels of sheltering were observed with these two cephalosporins, with an approximately two-fold increase in EC_50_ values occurring in both instances compared to the non-β-lactamase-producing control. The similar levels of sheltering provided by CTX-M-15 align with previous kinetic studies reporting similar *k*_cat_/*K*_M_ values for purified CTX-M-15 with cefazolin and ceftriaxone.^57^ However, the CTX-M-15-producing *E. coli* provided little-to-no sheltering against amoxicillin (Figure 5D, Table S1), consistent with the relatively low catalytic efficiency for CTX-M-15 against this penicillin.^58,59^

Our results have demonstrated the major role of β-lactamase substrate specificity in antibiotic sheltering. However, the rate of β-lactam hydrolysis that is catalysed by β-lactamases located in the periplasm of a Gram-negative bacterium will also depend on how easily a β-lactam antibiotic can enter the periplasm (Figure 1A).^60^ The passage of β-lactams across the Gram-negative OM often depends on porin proteins which form transmembrane channels.^61,62^ In *E. coli*, OmpC and OmpF are non-specific porins used by certain antibiotics to enter the cell.^63^ Gram-negative bacteria often become resistant to antibiotics through mutations to porin-encoding genes, reducing the entry of antibiotics such as β-lactams into the periplasm.^64^ To investigate whether changes to antibiotic entry impact β-lactamase-mediated antibiotic sheltering, we tested NDM-1-producing *E. coli* mutants in which the *ompF, ompC*, and *ompA* genes were disrupted.

In these sheltering assays, the NDM-1-producing Δ*ompC* and Δ*ompA E. coli* strains were observed to have greater EC_50_ values when compared to wild-type NDM-1-producing *E. coli* (Figure 6A). This suggests that these mutant strains in fact provide greater levels of sheltering compared to the wild-type strain. Consistent with this observation, both Δ*ompC* and Δ*ompA* strains degraded meropenem more quickly than the wild-type strain in a kinetic β-lactamase assay (Figure 6B).

**Figure 6.**
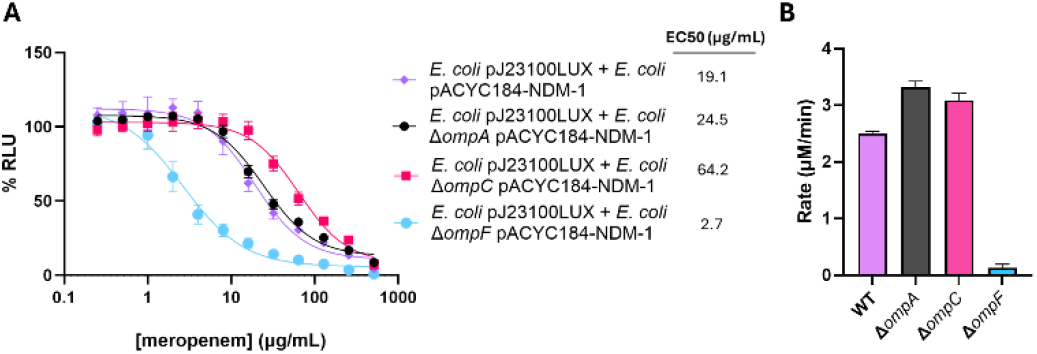
Meropenem sheltering and hydrolysis assays for NDM-1-producing *E. coli* BW25113 mutants lacking porins. A. Dose-response curves showing the EC_50_ values determined for the reporter strain in the presence of meropenem and NDM-1-producing *E. coli* WT, Δ*ompA*, Δ*ompC*, and Δ*ompF*. B. Initial velocities of meropenem (80 μM) hydrolysis by NDM-1-producing *E. coli* (WT and variants) as determined by UV-Vis spectrophotometry. 95% confidence intervals are reported in Table S1. % RLU = percent normalized relative luminescence units. All mutant strains were transformed with pACYC184-NDM-1. n = 3. Error bars represent standard deviations.

The porin OmpC has been shown to play an important role in the entry of carbapenems,^65^ and strains overexpressing OmpC exhibit lower minimum inhibitory concentrations (MICs) for carbapenems.^66^ While this would suggest that the Δ*ompC* strain should degrade meropenem more slowly, disruption of the *ompC* gene has been demonstrated to destabilize the OM.^61^ As a result, this likely leads to increased β-lactam entry and consequently greater antibiotic degradation and sheltering.

Researchers have observed that OmpA plays a major role in the stability of the *E. coli* cell wall by non-covalently anchoring the OM to peptidoglycan.^67^ Loss of OmpA compromises the OM, rendering bacteria more susceptible to stresses including antibiotics.^61,67^ Similar to what was seen with the Δ*ompC* strain, the loss of OmpA may allow more β-lactams into the periplasm, or could potentially increase the release of β-lactamases into the extracellular environment.

In contrast to the other two mutants, the Δ*ompF* strain provided far lower levels of sheltering with an EC_50_ 6-fold lower than that observed for wild-type NDM-1-producing *E. coli* (Figure 6A, Table S1). OmpF is used by carbapenems and other β-lactams to enter the periplasm,^68,69^ and disrupting the *ompF* gene increases the MICs of carbapenems.^70,71^ In our UV-Vis β-lactamase assays, the NDM-1-producing Δ*ompF E. coli* strain degraded meropenem much more slowly than the other *E. coli* strains tested (Figure 6B). With less meropenem reaching the periplasm of the NDM-1-producing bacteria, less antibiotic degradation occurs, exposing the susceptible reporter strain to higher antibiotic concentrations. Notably, the loss of OmpF is frequently observed in clinical isolates as an antibiotic resistance mechanism.^68^

Our findings show that antibiotic sheltering is influenced not only by the specific β-lactam-β-lactamase combination under investigation, but also by cellular factors that influence the ability of β-lactamases to interact with their targets. Beyond differences in porin expression, factors such as efflux pumps and lipopolysaccharide structure will also impact the accumulation of antibiotics in the periplasm. As such, the extent of sheltering provided by clinically relevant β-lactamase-producing bacteria will likely differ from what was observed with lab strains.

We next extended our sheltering assays to the study of three carbapenemase-producing clinical isolates (Figure 7). The VIM-producing *Enterobacter cloacae* strain provided the highest level of sheltering with an EC_50_ value of 19, while the KPC-2-producing *Klebsiella oxytoca* and NDM-producing *Klebsiella pneumoniae* strains had similar EC_50_ values of 10 and 11, respectively (Figure 7A, Table S1). In these experiments, the non-β-lactamase-producing *Escherichia coli* ATCC 25922 strain served as a negative control and offered no sheltering. The results obtained from these sheltering assays followed similar trends as seen through UV-Vis kinetic analyses of meropenem hydrolysis, where the *E. cloacae* strain demonstrated the fastest rate of hydrolysis, followed by the *K. pneumoniae* and *K. oxytoca* strains (Figure 7B). The results obtained from clinical isolates differ from those of the laboratory strains carrying the same β-lactamase, suggesting that variations in porin expression and other genetic mutations may influence bacterial sheltering. However, due to the absence of complete genome sequences for these isolates, it is not possible to determine the specific genetic factors contributing to the observed differences in sheltering.

**Figure 7.**
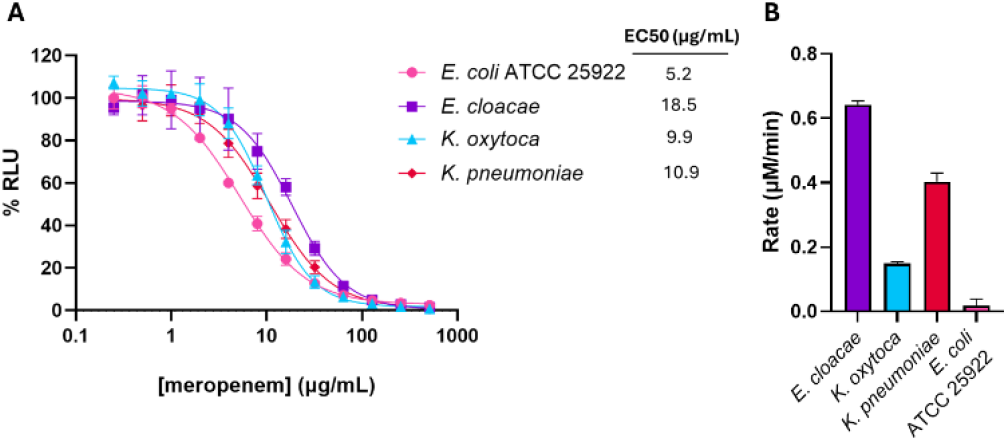
Meropenem sheltering and hydrolysis assays for β-lactamase-producing clinical isolates. A. Dose-response curves showing the EC_50_ values determined for the *E. coli* ATCC 25922 non-β-lactamase-producing control, a VIM-producing *E. cloacae* strain, a KPC-producing *K. oxytoca* strain, and an NDM-producing *K. pneumoniae* strain in the presence of meropenem. B. Initial velocities for meropenem (80 μM) hydrolysis by the clinical strains and *E. coli* ATCC 25922 as determined by UV-Vis spectrophotometry. 95% confidence intervals are reported in Table S1. % RLU = percent normalized relative luminescence units. n = 3. Error bars represent standard deviations.

Antibiotic sheltering is a phenomenon that should be considered when defining the treatment course for a bacterial infection, particularly if there is a polymicrobial etiology. The presence of multiple bacterial pathogens with different antibiotic resistance mechanisms can lead to treatment failure and severe complications.^12^ Furthermore, resistance genes can be harboured by commensal microbes,^72^ and the products of these microbes may also influence antibiotic treatment. Consequently, it is critical to recognize the impact that bacterial production of antibiotic-degrading enzymes can have on the survival of antibiotic-susceptible bacteria in the same environment. We show that, under the assay conditions tested, β-lactamase-producing bacteria can allow β-lactam-susceptible bacteria to survive antibiotic concentrations far greater than what would normally kill them.

The extent of sheltering is closely related to the rate of antibiotic degradation. Similar to previous reports,^9,26,43^ we observed that sheltering depends greatly on the particular combination of β-lactam and β-lactamase. Furthermore, the OM permeability of the β-lactamase-producing bacterium plays a major role in sheltering, influencing the extent to which β-lactams are exposed to β-lactamases. While destabilization of the OM can improve the survival of β-lactam-susceptible bacteria present in the same environment, the co-occurrence of multiple resistance mechanisms (e.g., β-lactamase production and loss of OmpF) can drastically decrease the extent of antibiotic sheltering that occurs. This is significant as clinical isolates often harbour multiple resistance mechanisms that can, in combination, greatly decrease their susceptibility to antibiotics.^73–75^

While our luminescence-based assay has proven effective for evaluating antibiotic sheltering, there are certain limitations. During our studies, we observed that increases in cell density reduced the luminescence measurements, presumably because the cells interfered with the ability of light to reach the detector. However, this limitation could be addressed by optimizing the incubation period to prevent overgrowth and ensure reliable luminescence measurements. Another limitation of this assay is that the *in vitro* conditions used do not reflect the environmental conditions occurring at the site of an infection (e.g., cell density, richness of growth medium). Nonetheless, our assay offers a fast and quantitative approach for investigating the factors that contribute to antibiotic sheltering. This strategy could be used to study interactions involving more than two bacterial strains, sheltering in the context of biofilms, or the sheltering provided by other antibiotic-degrading enzymes.^35^

Our studies focused on β-lactamases in the periplasm, but these enzymes can also be released into the extracellular environment as the result of cell lysis, or through packaging within outer membrane vesicles.^76^ Future studies will investigate how the localization of β-lactamases influences the extent of antibiotic sheltering. Additionally, further research will examine the impact of environmental factors on the bacterial cell wall, as well as the effects of other cell wall-targeting antibiotics.

## Conclusions

The presence of β-lactamase-producing bacteria can greatly enhance the survival of susceptible bacteria following their exposure to β-lactam antibiotics. The extent of sheltering depends on the identity of the β-lactam and β-lactamase, and also on the cell wall permeability of the β-lactamase-producing bacterium. Our findings emphasize the importance of evaluating the resistance mechanisms of all of the microbes present in the environment of an infection, particularly in the context of the treatment of polymicrobial infections.

## Materials and methods

### Reagents and materials

Sterile 2TY media (16 g/L tryptone, 10 g/L yeast extract, and 5 g/L sodium chloride) was used to culture bacteria, with tryptone and yeast extract purchased from BioShop and sodium chloride from Fisher Scientific. Meropenem trihydrate, imipenem, cefazolin, kanamycin monosulfate salt and ceftriaxone disodium salt hemi(heptahydrate) were purchased from Glentham Life Sciences. Chloramphenicol was purchased from Fisher Scientific.

### Bacterial strains and plasmids

The plasmid pJ23100LUX was prepared by cloning the *luxCDABE* operon (amplified from pAKgfplux2; Addgene #14083) between the NcoI and SacI restriction sites of pIDMv5K (provided by Sebastian Cocioba), placing the operon under the control of the strong constitutive J23100 promoter (Figure S1). *E. coli* BW25113 transformed with pJ23100LUX was used as the luminescent β-lactam-susceptible reporter strain. The gene and promoter for CTX-M-15 were amplified from the genomic DNA of a *Klebsiella pneumoniae* strain by polymerase chain reaction (PCR) and cloned into pACYC184 (National BioResource Project, NBRP) similar to our previous report.^77^ The gene and promoter for NDM-1 were amplified from pACYC184-NDM-1 by PCR and cloned into the EcoRI site of pHSG298 (NBRP) by in-cell ligation.^77^ *E. coli* BW25113 and single-gene knockout mutants from the Keio collection (Δ*ompA*, Δ*ompC*, and Δ*ompF*) were obtained from the NBRP.^78,79^ β-lactamase-producing lab strains were prepared by transforming *E. coli* BW25113 and the single gene knockout mutants with pACYC184 or pHSG298 plasmids encoding β-lactamases NDM-1, KPC-2, TEM-116, IMP-1, CTX-M-15, and OXA-48.^77^ Clinical isolates were provided by the clinical microbiology lab at Kingston General Hospital (KGH).

### Growth and antibiotic killing curves

The β-lactam-susceptible *E. coli* BW25113 pJ23100LUX reporter strain was grown overnight in 2TY media containing 50 µg/mL kanamycin at 37 °C and 200 rpm. A subculture was prepared by inoculating (1%) fresh 2TY (supplemented with 50 µg/mL kanamycin) with the overnight culture, which was then incubated at 37 °C and 200 rpm. The growth curve was determined by collecting luminescence and optical density (OD) measurements every hour using a Synergy LX multimode plate reader (Agilent BioTek). Clear non-treated Falcon 96-well microplates (Corning) were used for OD measurements. White LUMITRAC 200 96-well plate (Greiner Bio-One) were used for luminescence measurements. Killing curves were generated following the same protocol, except 2 µg/mL meropenem or imipenem was added to the subculture at time 0. Quadruplicates of each sample were measured.

### Enzyme kinetics

β-lactamase activity was measured by UV-Vis spectrophotometry. *E. coli* cells expressing NDM-1, KPC-2, IMP-1, OXA-48, TEM-116, or CTX-M-15 were cultured on 2TY agar plates supplemented with 25 µg/mL chloramphenicol. Suspensions of these *E. coli* cells were prepared in Dulbecco’s phosphate-buffered saline (DPBS) to an OD600 of 0.15. Stock solutions of meropenem and imipenem were prepared to a concentration of 80 µM using distilled water. In a 96-well UV-Star microplate (Greiner Bio-one), 100 µL of antibiotic solution and 100 µL of cell suspension were mixed and the change in absorbance at 297 nm over time was measured using a Synergy LX multimode plate reader (Agilent BioTek). Initial velocities were obtained by plotting antibiotic concentrations against time, and a linear regression fit was calculated using GraphPad Prism 10.2.3. Extinction coefficients of 11,500 M^−1^ cm^−1^ and 10,940 M^−1^ cm^−1^ were used for imipenem and meropenem, respectively.^52^

### Sheltering assays

Untransformed *E. coli* BW25113, the *E. coli* BW25113 pJ23100LUX luminescent reporter strain, the β-lactamase-producing *E. coli* BW25113 strains were cultured on 2TY agar (supplemented with kanamycin or chloramphenicol as needed) overnight at 37 °C. Unless otherwise stated, colonies were suspended in liquid 2TY to an OD600 of 0.1. Serial dilutions of the antibiotics tested were prepared in 2TY media. In a white LUMITRAC 200 96-well plate (Greiner Bio-One), 100 µL of antibiotic solution was mixed with a 100 µL suspension of a β-lactamase-producing strain and a 100 µL suspension of the reporter strain. The plate was incubated at 37 °C and luminescence was measured using a Synergy LX multimode plate reader after one hour. Control samples without β-lactamase-producing bacteria consisted of 100 μL antibiotic, 100 μL of the reporter strain suspension, and a 100 μL suspension of untransformed *E. coli* BW25113 cells. Luminescence values were blanked and normalized within each data set and plotted against antibiotic concentration. Dose-response curves were obtained using a non-linear fit analysis in GraphPad Prism 10.2.3 for reporter cells with β-lactamase producing cells (sheltered), and for reporter cells with untransformed cells (not sheltered).

## Data availability

The data supporting this article have been included as part of the Supplementary Information.

## Notes and references

1 A. R. Pacheco and D. Segrè, FEMS Microbiol. Lett., 2019, 366, 125.

2 K. A. Brogden, J. M. Guthmiller and C. E. Taylor, Lancet, 2005, 365, 3.

3 C. D. Sibley, H. Rabin and M. G. Surette, Future Microbiol., 2006, 1, 53–61.

4 M. D. Parkins and R. A. Floto, J. Cyst. Fibros., 2015, 14, 293– 304.

5 B. M. Peters, M. A. Jabra-Rizk, G. A. O’May, J. W. Costerton and M. E. Shirtliff, Clin. Microbiol. Rev., 2012, 25, 193–213.

6 L. M. Filkins and G. A. O’Toole, PLOS Pathog., 2015, 11, e1005258.

7 A. Hector, T. Kirn, A. Ralhan, U. Graepler-Mainka, S. Berenbrinker, J. Riethmueller, M. Hogardt, M. Wagner, A. Pfleger, I. Autenrieth, M. Kappler, M. Griese, E. Eber, P. Martus and D. Hartl, J. Cyst. Fibros., 2016, 15, 340–349.

8 A. Sandri, J. A. J. Haagensen, L. Veschetti, H. K. Johansen, S. Molin, G. Malerba, C. Signoretto, M. Boaretti and M. M. Lleo, Pathogens, 2021, 10, 978.

9 N. M. Smith, A. Ang, F. Tan, K. Macias, S. James, J. Sidhu and J. R. Lenhard, Antimicrob. Agents Chemother., 2021, 65, e02414–20.

10 G. Orazi and G. A. O’Toole, mBio, 2017, 8, e00873–17.

11 L. Radlinski, S. E. Rowe, L. B. Kartchner, R. Maile, B. A. Cairns, N. P. Vitko, C. J. Gode, A. M. Lachiewicz, M. C. Wolfgang and B. P. Conlon, PLOS Biol., 2017, 15, e2003981.

12 I. Brook, Clin. Microbiol. Infect., 2004, 10, 777–784.

13 M. Mora-Ochomogo and C. T. Lohans, RSC Med. Chem., 2021, 12, 1623–1639.

14 K. Bush and P. A. Bradford, Cold Spring Harb. Perspect. Med., 2016, 6, a025247.

15 C. L. Tooke, P. Hinchliffe, E. C. Bragginton, C. K. Colenso, V. H. A. Hirvonen, Y. Takebayashi and J. Spencer, J. Mol. Biol., 2019, 431, 3472–3500.

16 J. D. Pitout, C. C. Sanders and W. E. J. Sanders, Am. J. Med., 1997, 103, 51–59.

17 R. B. Sykes and M. Matthew, J. Antimicrob. Chemother., 1976, 2, 115–157.

18 T. Naas, S. Oueslati, R. A. Bonnin, M. L. Dabos, A. Zavala, L. Dortet, P. Retailleau and B. I. Iorga, J. Enzyme Inhib. Med. Chem., 2017, 32, 917–919.

19 K. Bush and P. A. Bradford, Clin. Microbiol. Rev., 2020, 33, e00047–19.

20 K. Bush, Antimicrob. Agents Chemother., 2018, 62, e01076–18.

21 P. Nordmann, G. Cuzon and T. Naas, Lancet Infect. Dis., 2009, 9, 228–236.

22 K. M. Papp-Wallace, A. Endimiani, M. A. Taracila and R. A. Bonomo, Antimicrob. Agents Chemother., 2011, 55, 4943– 4960.

23 J. Vergalli, I. V. Bodrenko, M. Masi, L. Moynié, S. Acosta-Gutiérrez, J. H. Naismith, A. Davin-Regli, M. Ceccarelli, B. Van Den Berg, M. Winterhalter and J.-M. Pagès, Nat. Rev. Microbiol., 2020, 18, 164–176.

24 S. W. Kim, J. S. Lee, S. B. Park, A. R. Lee, J. W. Jung, J. H. Chun, J. M. S. Lazarte, J. Kim, J.-S. Seo, J.-H. Kim, J.-W. Song, M. W. Ha, K. D. Thompson, C.-R. Lee, M. Jung and T. S. Jung, Int. J. Mol. Sci., 2020, 21, 2822.

25 M. Pavlaki, G. Poulakou, P. Drimousis, G. Adamis, E. Apostolidou, N. K. Gatselis, I. Kritselis, A. Mega, V. Mylona, A. Papatsoris, A. Pappas, A. Prekates, M. Raftogiannis, K. Rigaki, K. Sereti, D. Sinapidis, I. Tsangaris, V. Tzanetakou, D. Veldekis, K. Mandragos, H. Giamarellou and G. Dimopoulos, J. Glob. Antimicrob. Resist., 2013, 1, 207–212.

26 M. H. Perlin, D. R. Clark, C. McKenzie, H. Patel, N. Jackson, C. Kormanik, C. Powell, A. Bajorek, D. A. Myers, L. A. Dugatkin and R. M. Atlas, Proc. R. Soc. B Biol. Sci., 2009, 276, 3759–3768.

27 S. Caprari, 2021, 292.

28 Y.-T. Liao, S.-C. Kuo, M.-H. Chiang, Y.-T. Lee, W.-C. Sung, Y.-H. Chen, T.-L. Chen and C.-P. Fung, Antimicrob. Agents Chemother., 2015, 59, 7346–7354.

29 V. Schaar, T. Nordström, M. Mörgelin and K. Riesbeck, Antimicrob. Agents Chemother., 2011, 55, 3845–3853.

30 D. Hubert, H. Réglier-Poupet, I. Sermet-Gaudelus, A. Ferroni, M. Le Bourgeois, P.-R. Burgel, R. Serreau, D. Dusser, C. Poyart and J. Coste, J. Cyst. Fibros., 2013, 12, 497–503.

31 M. L. Maliniak, A. A. Stecenko and N. A. McCarty, J. Cyst. Fibros., 2016, 15, 350–356.

32 M. Bielaszewska, O. Daniel, O. Nyč and A. Mellmann, Membranes, 2021, 11, 806.

33 R. Cantón, A. Novais, A. Valverde, E. Machado, L. Peixe, F. Baquero and T. M. Coque, Clin. Microbiol. Infect., 2008, 14, 144–153.

34 A. P. Johnson and N. Woodford, J. Med. Microbiol., 2013, 62, 499–513.

35 R. A. Sorg, L. Lin, G. S. van Doorn, M. Sorg, J. Olson, V. Nizet and J.-W. Veening, PLOS Biol., 2016, 14, e2000631.

36 G. Wright, Adv. Drug Deliv. Rev., 2005, 57, 1451–1470.

37 N. M. Vega and J. Gore, Curr. Opin. Microbiol., 2014, 21, 28–34.

38 L. Geyrhofer, P. Ruelens, A. D. Farr, D. Pesce, J. A. G. M. de Visser and N. Brenner, mBio, 2023, e02456–22.

39 D. Clark R., Front. Biosci., 2009, 14, 4815.

40 E. A. Yurtsev, H. X. Chao, M. S. Datta, T. Artemova and J. Gore, Mol. Syst. Biol., 2013, 9, 683.

41 L. A. Dugatkin, M. Perlin, J. S. Lucas and R. Atlas, Proc. R. Soc. B Biol. Sci., 2005, 272, 79–83.

42 F. Medaney, T. Dimitriu, R. J. Ellis and B. Raymond, ISME J., 2016, 10, 778–787.

43 Y.-T. Liao, S.-C. Kuo, Y.-T. Lee, C.-P. Chen, S.-W. Lin, L.-J. Shen, C.-P. Fung, W.-L. Cho and T.-L. Chen, Antimicrob. Agents Chemother., 2014, 58, 3983–3990.

44 G. Gazzola, O. Habimana, L. Quinn, E. Casey and C. D. Murphy, Biofouling, 2019, 35, 299–307.

45 J. Chiou, T. Y.-C. Leung and S. Chen, Antimicrob. Agents Chemother., 2014, 58, 5372–5378.

46 G. Cuzon, T. Naas, P. Bogaerts, Y. Glupczynski, T.-D. Huang and P. Nordmann, Antimicrob. Agents Chemother., 2008, 52, 3463–3464.

47 T.-L. Lin, S.-I. Tang, C.-T. Fang, P.-R. Hsueh, S.-C. Chang and J.-T. Wang, Microb. Drug Resist., 2006, 12, 12–15.

48 F. Fonseca, E. I. Chudyk, M. W. van der Kamp, A. Correia, A. J. Mulholland and J. Spencer, J. Am. Chem. Soc., 2012, 134, 18275–18285.

49 M. I. El-Gamal, I. Brahim, N. Hisham, R. Aladdin, H. Mohammed and A. Bahaaeldin, Eur. J. Med. Chem., 2017, 131, 185–195.

50 B. A. Lund, A. M. Thomassen, T. J. W. Carlsen and H.-K. S. Leiros, Acta Crystallogr. Sect. F Struct. Biol. Commun., 2021, 77, 312–318.

51 B. A. Lund, T. Christopeit, Y. Guttormsen, A. Bayer and H.-K. S. Leiros, J. Med. Chem., 2016, 59, 5542–5554.

52 A. M. Queenan, W. Shang, R. Flamm and K. Bush, Antimicrob. Agents Chemother., 2010, 54, 565–569.

53 V. Stojanoski, L. Hu, B. Sankaran, F. Wang, P. Tao, B. V. V. Prasad and T. Palzkill, ACS Infect. Dis., 2021, 7, 445–460.

54 D. Y. Wang, M. I. Abboud, M. S. Markoulides, J. Brem and C. J. Schofield, Future Med. Chem., 2016, 8, 1063–1084.

55 Y. Pfeifer, A. Cullik and W. Witte, Int. J. Med. Microbiol., 2010, 300, 371–379.

56 Bradford Patricia A., Clin. Microbiol. Rev., 2001, 14, 933–951.

57 M. Faheem, M. T. Rehman, M. Danishuddin and A. U. Khan, PLoS ONE, 2013, 8, e56926.

58 L. Poirel, J. Antimicrob. Chemother., 2002, 50, 1031–1034.

59 L. Poirel, J.-M. O. De La Rosa, A. Richard, M. Aires-de-Sousa and P. Nordmann, Antimicrob. Agents Chemother., 2019, 63, e01515–19.

60 I. Massova and S. Mobashery, Antimicrob Agents Chemother, 1998, 42, 17.

61 U. Choi and C.-R. Lee, Front. Microbiol., 2019, 10, 953.

62 J. D. Prajapati, U. Kleinekathöfer and M. Winterhalter, Chem. Rev., 2021, 121, 5158–5192.

63 S. W. Cowan, T. Schirmer, G. Rummel, M. Steierf, R. Ghosh, R. A. Pauptitt, J. N. Jansonius and J. P. Rosenbusch,.

64 A. H. Delcour, Biochim. Biophys. Acta BBA – Proteins Proteomics, 2009, 1794, 808–816.

65 H. Lou, M. Chen, S. S. Black, S. R. Bushell, M. Ceccarelli, T. Mach, K. Beis, A. S. Low, V. A. Bamford, I. R. Booth, H. Bayley and J. H. Naismith, PLoS ONE, 2011, 6, e25825.

66 Q.-T. Tran, R. A. Pearlstein, S. Williams, J. Reilly, T. Krucker and G. Erdemli, Proteins Struct. Funct. Bioinforma., 2014, 82, 2998–3012.

67 Y. Wang, Biochem. Biophys. Res. Commun., 2002, 292, 396–401.

68 H. Yigit, A. M. Queenan, J. K. Rasheed, J. W. Biddle, A. Domenech-Sanchez, S. Alberti, K. Bush and F. C. Tenover, Antimicrob Agents Chemother.

69 R. E. Hancock, J. Bacteriol., 1987, 169, 929–933.

70 K. J. Harder, H. Nikaido and M. Matsuhashi, Antimicrob. Agents Chemother., 1981, 20, 549–552.

71 B. K. Ziervogel and B. Roux, Structure, 2013, 21, 76–87.

72 T. S. Ghosh, S. S. Gupta, G. B. Nair and S. S. Mande, PLOS ONE, 2014, 8, e83823.

73 R. Ferrer-Espada, S. Sánchez-Gómez, B. Pitts, P. S. Stewart and G. Martínez-de-Tejada, Int. J. Antimicrob. Agents, 2020, 56, 105986.

74 P. M. Hawkey and C. J. Munday, Rev. Med. Microbiol., 2004, 15, 51–61.

75 É. Ruppé, P.-L. Woerther and F. Barbier, Ann. Intensive Care, 2015, 5, 21.

76 M. M. B. Martínez, R. A. Bonomo, A. J. Vila, P.C. Maffía and L. J. González, mBio, 2021, 12, e01836–21.

77 M. A. Jeffs, R. A. V. Gray, P. M. Sheth and C. T. Lohans, Chem. Commun., 2023, 59, 12707–12710.

78 T. Baba, T. Ara, M. Hasegawa, Y. Takai, Y. Okumura, M. Baba, K. A. Datsenko, M. Tomita, B. L. Wanner and H. Mori, Mol. Syst. Biol., DOI:10.1038/msb4100050.

79 Y. Yamazaki, R. Akashi, Y. Banno, T. Endo, H. Ezura, K. Fukami-Kobayashi, K. Inaba, T. Isa, K. Kamei, F. Kasai, M. Kobayashi, N. Kurata, M. Kusaba, T. Matuzawa, S. Mitani, T. Nakamura, Y. Nakamura, N. Nakatsuji, K. Naruse, H. Niki, E. Nitasaka, Y. Obata, H. Okamoto, M. Okuma, K. Sato, T. Serikawa, T. Shiroishi, H. Sugawara, H. Urushibara, M. Yamamoto, Y. Yaoita, A. Yoshiki and Y. Kohara, Nucleic Acids Res., 2010, 38, D26–D32.

