## Supplemental Information for "Contributions of β-lactamase substrate specificity and outer membrane permeability to the antibiotic sheltering of β-lactam-susceptible bacteria"

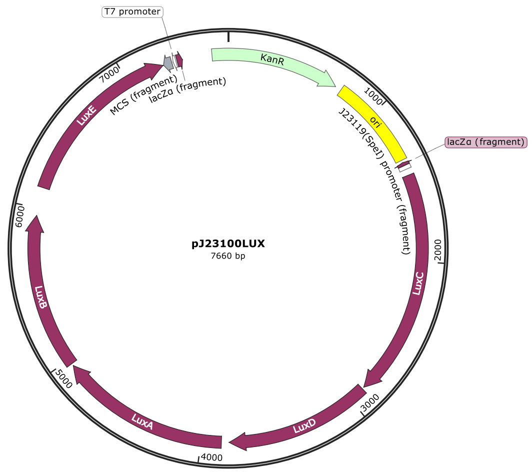


**Figure S1. pJ23100LUX plasmid map.** Whole plasmid sequencing was performed by Plasmidsaurus using Oxford Nanopore Technology with custom analysis and annotation. The *lux* operon (*luxCDABE*) is under the control of the strong constitutive J23100 promoter. This plasmid also carries a kanamycin resistance gene.

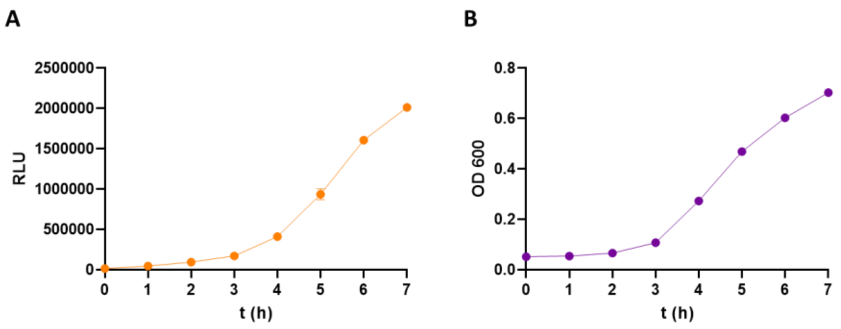


**Figure S2. Initial characterization of the *Escherichia coli* BW25113 pJ23100LUX reporter strain.** **(A)** Luminescence measurements monitoring the growth of *E. coli* pJ23100LUX over time, represented in relative luminescence units (RLU). **(B)** Growth of *E. coli* pJ23100LUX cells over time as measured by optical density (OD) at 600 nm.

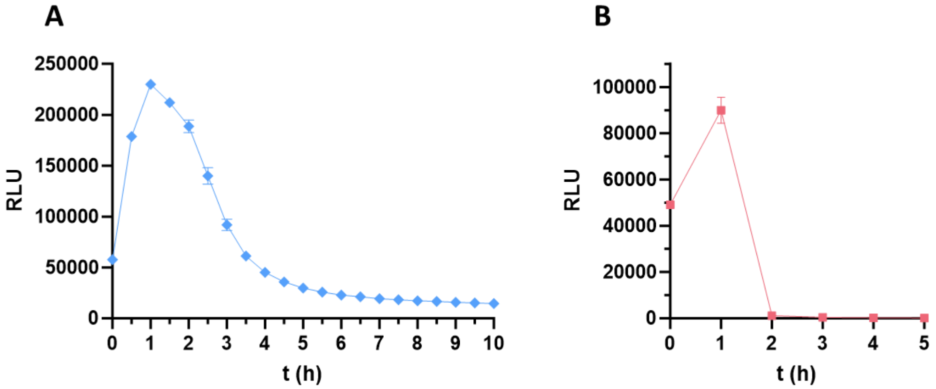


**Figure S3. Killing curves showing the impact of carbapenems on the reporter strain.** *E. coli* BW25113 pJ23100LUX cells were treated with 2TY media supplemented with **(A)** 2 μg/mL meropenem, or **(B)** 2 μg/mL imipenem. RLU: Relative luminescence units. n = 3. Error bars represent standard deviations.

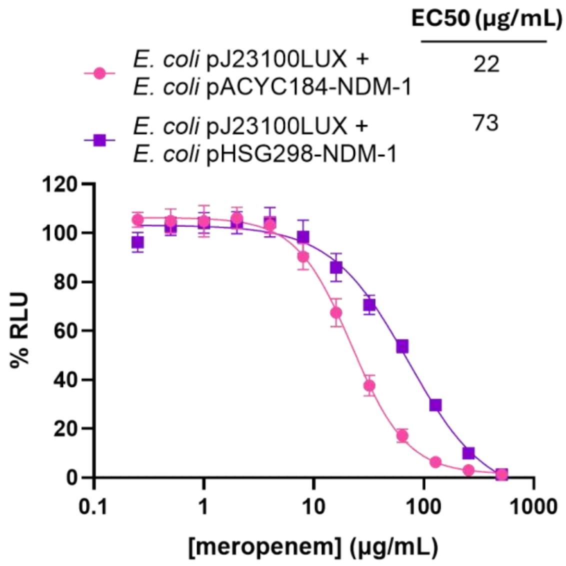


**Figure S4. Sheltering of the luminescent reporter strain by *E. coli* transformed with pHSG298-NDM-1 or pACYC184-NDM-1.** Dose-response curves showing the $EC_{50}$ values for both high-copy (pHSG298) and low-copy (pACYC184) plasmids that encode for the β-lactamase NDM-1. The strain transformed with the pHSG298 plasmid provides greater levels of sheltering under the assay conditions. % RLU = percent normalized relative luminescence percentage. n = 4. Error bars represent standard deviations.

**Table S1.** $EC_{50}$ values and 95% confidence intervals from dose-response curves.

| Figure No. | Sheltering strain | β-lactam | $EC_{50}$ (95% Confidence Interval) |
|---|---|---|---|
| 2B | *E. coli* pACYC184-NDM-1 (OD 0.01) | Meropenem | 5.5 (4.8 - 6.5) |
| | *E. coli* pACYC184-NDM-1 (OD 0.05) | | 9.7 (8.4 - 11.6) |
| | *E. coli* pACYC184-NDM-1 (OD 0.1) | | 23.6 (19.3 - 29.7) |
| | *E. coli* WT (OD 0.1) | | 4.4 (3.5 - 5.9) |
| 3A | *E. coli* pACYC184-NDM-1 | Meropenem | 23.6 (21.9 - 25.6) |
| | *E. coli* WT | | 3.6 (3.1 - 4.4) |
| 3B | *E. coli* pACYC184-KPC-2 | Meropenem | 18.0 (17.2 - 18.9) |
| | *E. coli* WT | | 2.4 (2.2 - 2.6) |
| 3C | *E. coli* pACYC184-IMP-1 | Meropenem | 21.3 (20.4 - 22.3) |
| | *E. coli* WT | | 1.9 (1.8 - 2.1) |
| 3D | *E. coli* pACYC184-OXA-48 | Meropenem | 2.5 (2.2 - 2.7) |
| | *E. coli* WT | | 1.9 (1.8 - 2.1) |
| 3E | *E. coli* pACYC184-TEM-116 | Meropenem | 3.0 (2.9 - 3.1) |
| | *E. coli* WT | | 3.1 (2.9 - 3.5) |
| 4A | *E. coli* pACYC184-NDM-1 | Imipenem | 23.6 (21.1 - 26.3) |
| | *E. coli* WT | | 1.1 (1.0 - 1.3) |
| 4B | *E. coli* pACYC184-KPC-2 | Imipenem | 28.4 (26.0 - 31.1) |
| | *E. coli* WT | | 4.4 (4.0 - 4.9) |
| 4C | *E. coli* pACYC184-OXA-48 | Imipenem | 3.6 (lower limit 3.4)[a] |
| | *E. coli* WT | | 1.1 (1.0 - 1.2) |
| 5A | *E. coli* pACYC184-KPC-2 | Meropenem + Avibactam | 4.7 (4.0 - 5.5) |
| | *E. coli* WT | | 5.0 (4.2 - 6.0) |
| 5B | *E. coli* pACYC184-CTX-M-15 | Cefazolin | 30.0 (28.7- 30.7) |
| | *E. coli* WT | | 17.4 (16.8 - 18.1) |
| 5C | *E. coli* pACYC184-CTX-M-15 | Ceftriaxone | 43.9 (40.4 - 47.7) |
| | *E. coli* WT | | 21.6 (17.4 - 26.6) |
| 5D | *E. coli* pACYC184-CTX-M-15 | Amoxicillin | 23.7 (23.0 - 24.4) |
| | *E. coli* WT | | 21.4 (20.7 - 22.1) |
| 6A | *E. coli* Δ*ompA* pACYC184-NDM-1 | Meropenem | 24.5 (21.6 - 28.1) |
| | *E. coli* Δ*ompC* pACYC184-NDM-1 | | 64.2 (53.0 - 81.6) |
| | *E. coli* Δ*ompF* pACYC184-NDM-1 | | 2.7 (2.1 - 3.2) |
| | *E. coli* pACYC184-NDM-1 | | 19.1 (15.9 - 23.3) |
| 7A | *E. coli* ATCC 25922 | Meropenem | 5.2 (4.8 - 5.7) |
| | *Enterobacter cloacae* clinical isolate | | 18.5 (15.1 - 22.7) |

| | | | |
|---|---|---|---|
| | *Klebsiella oxytoca* clinical isolate | | 10.0 (9.1 - 10.9) |
| | *Klebsiella pneumoniae* clinical isolate | | 10.9 (9.5 - 12.4) |
| S4 | *E. coli* pHSG298-NDM-1 | Meropenem | 22.2 (20.5 - 23.9) |
| | *E. coli* pACYC184-NDM-1 | | 72.5 (60.9 - 89.9) |

[a] GraphPad Prism 10.2.3 was unable to calculate a complete confidence interval.
